# An Interpretable Graph-Regularized Optimal Transport Framework for Diagonal Single-Cell Integrative Analysis

**DOI:** 10.1101/2024.10.30.621072

**Authors:** Zexuan Wang, Qipeng Zhan, Shu Yang, Zhuoping Zhou, Mengyuan Kan, Tianhuan Zhai, Li Shen

## Abstract

**Background:** Recent advancements in single-cell omics technologies have enabled detailed characterization of cellular processes. However, coassay sequencing technologies remain limited, resulting in un-paired single-cell omics datasets with differing feature dimensions;

**Finding:** we present GROTIA (Graph-Regularized Optimal Transport Framework for Diagonal Single-Cell Integrative Analysis), a computational method to align multi-omics datasets without requiring any prior correspondence information. GROTIA achieves global alignment through optimal transport while preserving local relationships via graph regularization. Additionally, our approach provides interpretability by deriving domain-specific feature importance from partial derivatives, highlighting key biological markers. Moreover, the transport plan between modalities can be leveraged for post-integration clustering, enabling a data-driven approach to discover novel cell subpopulations;

**Conclusions:** We demonstrate GROTIA’s superior performance on four simulated and four real-world datasets, surpassing state-of-the-art unsupervised alignment methods and confirming the biological significance of the top features identified in each domain. The software is available at https://github.com/PennShenLab/GROTIA.

## Introduction

The advancement of single-cell technology offers a comprehensive understanding of cellular heterogeneity and the dynamic evolution of cell states. Various single-cell measurements reveal different aspects: scRNA-seq [1, 2] provides detailed gene expression profiles, while scATAC-seq [3] sheds light on chromatin accessibility in individual cells. Integrating these datasets is crucial as it allows for a more holistic view of cellular mechanisms, enabling the correlation of transcriptional activity with chromatin states to better understand gene regulation and cellular function.

Lots of computational methods have recently been developed to integrate data across multiple modalities [4, 5]. A critical challenge for these algorithms is their reliance on correspondence information to identify alignments between paired cells. In practice, such information is often only partially available, hindering the effectiveness of existing strategies [6, 7, 8]. This limitation has led researchers to focus on diagonal integration under semi-supervised settings, where alignment is achieved without direct cell-to-cell correspondences, but cell type labels are still used for hyperparameter tuning. The Generalized Unsupervised Manifold Alignment (GUMA) method [9] aligns datasets by optimizing local geometric structures to establish a one-to-one correspondence. Building upon this, the Unsupervised topological alignment for single-cell multi-omics integration (UnionCom) algorithm by Cao et al. [10] enables semi-supervised topological alignment, relaxing GUMA’s strict one-to-one mapping requirement. Liu et al. [11] proposed an alternative manifold alignment strategy called MMD-MA, which employs the Maximum Mean Discrepancy (MMD) metric for alignment. Additionally, the Single-Cell Multi-Omics Alignment with Optimal Transport (SCOT) method [12] utilizes Gromov-Wasserstein distances to align multi-omic single-cell data. Autoencoder based method have also been proposed to align data across modality to use per ae per modality and align them in shared latent space [13, 14]. However, even when integration is performed without correspondences, hyperparameters are often tuned using cell label validation, rendering the process semi-supervised. Demetci et al. [12] demonstrated that most methods fail to adapt to fully unsupervised settings when no orthogonal alignment information is available.

Diagonal single-cell multi-omics integration thus faces several key obstacles. First, in the absence of paired samples, one must work under an unpaired assumption, which is common given the practical and financial difficulties of obtaining perfectly matched datasets. Second, integration often relies on shared features (e.g., overlapping genes) that may be missing or poorly represented across modalities. Third, most computational pipelines rely on label-based metrics for hyperparameter tuning, which is problematic in truly unsupervised settings where no external annotations exist. Finally, many existing methods lack an interpretable framework to explain the learned shared embeddings.

Here, we propose GROTIA, a fully unsupervised diagonal integration method that uses optimal transport and graph regularization to establish alignment without relying on one-to-one correspondences or labeled data. We embed each dataset in a high-dimensional kernel space to capture cell– cell similarities, then learn mappings that transform each dataset into a shared lower-dimensional space for direct comparison. Our framework preserves local geometry via graph Laplacian regularization while performing global alignment through optimal transport, thereby avoiding the need for label-based hyperparameter tuning. In addition, we provide gradient-based sensitivity analyses to highlight key biological genes and peaks that drive the alignment, clear interpretability of the learned latent representation.

We extensively evaluate our model against SCOT, MMD-MA, UnionCom, Uniport, Scconfluence across four simulated and four real-world datasets in unsupervised and semi-supervised settings. Our Graph-Regularized Optimal Transport (GROTIA) algorithm matches the performance of state-of-the-art methods. The schematic design of our approach is illustrated in **Figure 1**. The source code for GROTIA is publicly available at https://github.com/PennShenLab/GROTIA.

**Figure 1.**
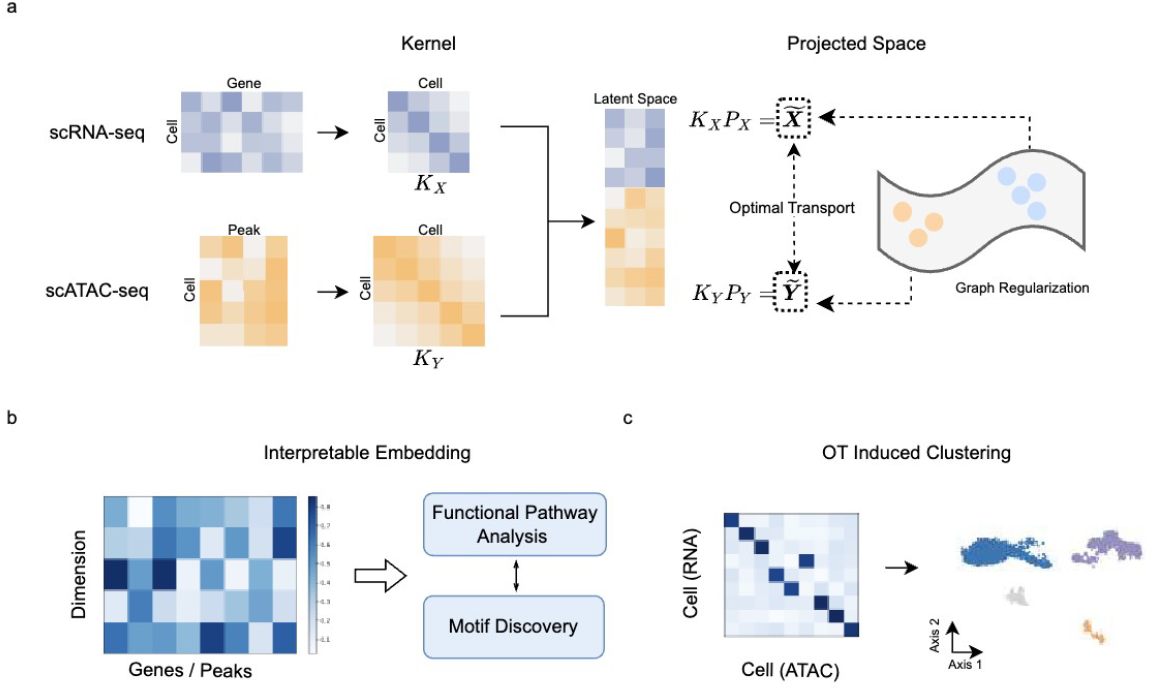
Overview of the GROTIA Framework for Multi-Omics Single-Cell Integration. (a) Schematic Design: For each single-cell modality (e.g., scRNA-seq, scATAC-seq), GROTIA constructs a kernel matrix capturing pairwise cell similarities (e.g., *K*_*X*_ and *K*_*Y*_). It then learns mapping matrices *P*_*X*_ and *P*_*Y*_ to project cells from each modality’s RKHS into a shared latent space, where distributions are aligned via optimal transport. A graph regularization term preserves local neighborhood structure, ensuring that cells close in the original domain remain similarly positioned after integration. (b) Interpretable Embedding: Once the shared embeddingis obtained, GROTIA provides dimensionwise importance scores for genes or peaks. These scores can be used for downstream analyses such as Gene Ontology (GO) or motif discovery, provide biological interpretation of each latent dimension. (c) OT-Induced Co-Clustering: GROTIA leverages the cross-modality transport plan, which quantifies how strongly each scRNA cell aligns with each scATAC cell. By simultaneously grouping cells from both modalities according to these alignment strengths, GROTIA identifies co-clusters of subpopulations with closely matched regulatory states in the latent space.

## Methods

### Simulated Datasets

We evaluated the GROTIA algorithm using four simulated datasets: three from Liu et al. [11], specifically designed to test alignment methods with different geometric structures, and one additional dataset from Demetci et al. [12], simulating single-cell RNA sequencing count data via Splatter [15]. Specifically, the first dataset presents a branch structure in two-dimensional space, the second a Swiss roll in three-dimensional space, and the third a circular frustum also in three-dimensional space. Although these datasets originally feature complex topological and geometric structures in low-dimensional spaces, they have been nonlinearly projected into high-dimensional feature spaces of 1000 and 2000 dimensions for evaluating alignment methods. The fourth is a synthetic RNA-seq dataset from Demetci et al. [12], consisting of 5,000 cells with either 50 or 500 features. Following the approaches in the original publications, we applied Z-score normalization to all features before running alignment algorithms.

### Real World Datasets

We then evaluated the GROTIA algorithm on four real-world datasets, widely recognized as gold-standard benchmarks in multi-omics integration and commonly used for assessing state-of-the-art methods. These include: (1) scGEM, which simultaneously profiles gene expression and DNA methylation [16].; (2) a dataset generated by the SNARE-seq assay, linking chromatin accessibility with gene expression [17].; (3) a human PBMC dataset (PBMC 10X) consisting of 9,378 cells per modality [14].; and (4) an additional PBMC 1OX dataset containing 11,259 cells per modality [13]. We chose these paired multi-omics datasets specifically to enable diagonal integration with known ground-truth cell correspondences. Importantly, during benchmarking, all methods are provided with unpaired data, and the known cell-pairing information was used only for evaluating alignment accuracy.

The first real-world dataset, named “scGEM,” measures gene expression and DNA methylation in the same cells and was generated using the scGEM assay. It contains human somatic cells reprogrammed to a pluripotent state, forming a continuous developmental trajectory. Cao et al. [10] and Demetci et al. [12] previously employed this dataset to evaluate integration methods. Specifically, it has 177 cells with 34 gene-expression features and 177 cells with 27 DNA methylation features. We used the preprocessed version from Demetci et al. [12].

The second real-world dataset, SNARE-seq, jointly profiles chromatin accessibility and gene expression. The dataset was pre-processed using cisTopic [18], resulting in an ATAC-seq matrix of 1,047 cells by 19 features and an RNA-seq matrix of 1,047 cells by 10 features. Following standard practice, unit normalization was then applied to these matrices. We used this preprocessed SNARE-seq dataset from Demetci et al. [12].

Additionally, we analyzed two multi-omics peripheral blood mononuclear cell (PBMC) datasets from publicly available 10x Genomics sources. The first, preprocessed by UniPort [14], contains 11,259 cells with 28,307 scATAC-seq features and 11,942 scRNA-seq genes. The second, preprocessed by scConfluence [13], includes 9,378 cells with 130,417 scATAC-seq features and 15,417 scRNA-seq genes. We used both PBMC datasets as provided for our integrative analyses.

### Problem Formulation

We introduce a method to integrate single-cell datasets across different conditions or modalities. Let us consider two datasets, *X* and *Y*, with respective representations 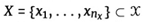 and 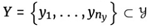, where *n*_*x*_ and *n*_*y*_ denote the number cells in *X* and *Y*, respectively. We aim to uncover a shared manifold structure between *X* and *Y* without a priori correspondence between the datasets.

To achieve this, we first compute the intra-dataset kernels *K*_*X*_ and *K*_*Y*_, which capture the internal structure of *X* and *Y*, respectively. As long as it is positive definite, each kernel corresponds to an implicit feature mapping ϕ_*X*_: 𝒳 → ℋ_*X*_ and ϕ_*Y*_: *y* → ℋ_*Y*_, where ℋ_*X*_ and 𝒦_*Y*_ are the Reproducing Kernel Hilbert Spaces (RKHS) associated with *K*_*X*_ and *K*_*Y*_. Subsequently, we seek mapping functions *f*_*X*_ : 𝒳 → *R*^*k*^ and *f*_*Y*_ : *y* → *R*^*k*^, where *k* is the dimensionality of the shared space. These functions are optimized so that the mapped representations *f*_*X*_(*X*) andfy *f*_*Y*_*(Y)* are well-aligned, thereby discovering the shared manifold structure.

### Kernel Representation

To capture the intrinsic geometry of the datasets, we define the intra-dataset kernels *K*_*X*_ and *K*_*Y*_ using Gaussian kernel functions:

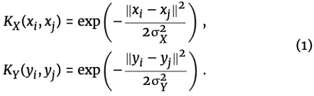

where *σ*_*X*_ and *σ*_*Y*_ are bandwidth parameters specific to *X* and *Y*, respectively. These kernels define the feature maps ϕ_*X*_: 𝒳 → ℋ_*X*_ and ϕ_*Y*_: *y* → ℋ_*Y*_ into their respective Reproducing Kernel Hilbert Spaces (RKHS).

We adopt a data-driven approach to determine *σ*_*X*_ and *σ*_*Y*_ by setting each parameter to the mean of the pairwise Euclidean distances within the corresponding dataset. This heuristic adjusts the bandwidths to reflect the average spatial dispersion of the data points, thereby tuning the kernels to the specific scale of variability in each dataset.

### Optimal Transport

For simplicity, we will use the notation: 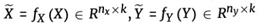 to represent the mapped datasets. The Sinkhorn divergence between the projected representations 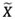 and 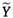 is defined as:

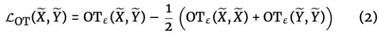

where OT_ε_ (·, ·) denotes the entropically regularized optimal transport cost between two distributions. Next, we define the entropic optimal transport cost between 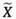 and 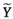. The cost is computed as:

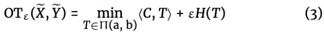

where 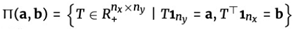. The matrix 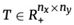 is the transport plan matrix, representing the amount of mass transported from 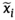 to 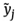.The cost matrix 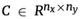 quantifies the pairwise distances between the projected samples. Each element is defined as: 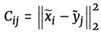. The parameter, *ε* > 0 is the entropic regularization parameter that smooths the optimization problem and 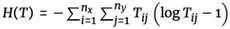 is the entropy of the transport plan *T*. The marginal distributions 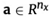 and 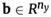 are typically uniform distributions over the samples: 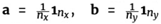, where 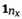, and 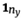 are vectors of ones with lengths *n*_*x*_ and *n*_*y*_, respectively. We utilize optimal transport over maximum mean discrepancy due to its benefits, such as non-vanishing gradients and other theoretical advantages [19].

### Graph Laplacian Regularization

To capture the local geometric structures of the datasets *X* and *Y*, we construct graph Laplacians based on the k-nearest neighbor relationships defined through the Gaussian kernels. These Laplacians serve as regularizers for the mapping functions, enforcing smoothness by ensuring that nearby data points in the RKHS spaces ℋ_*X*_ and ℋ_*Y*_ remain close in the shared latent space.

For each dataset, we begin by constructing a *k*-nearest neighbor graph in the RKHS. Specifically, for dataset *X*, we identify the set of *k*-nearest neighbors for each feature-mapped data point ϕ_*X*_ (*x*_*i*_), denoted as 𝒩_*k*_ ϕ_*X*_ (*x*_*i*_)), based on the distance metric in ℋ_*X*_. The adjacency matrix 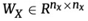 is then defined with entries:

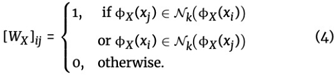

Similarly,for dataset Y, we construct the adjacency matrix 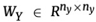. Next, we compute the degree matrices *D*_*X*_ and *D*_*Y*_, which are diagonal matrices where each diagonal entry represents the sum of the edge weights connected to a node: 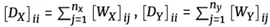. The graph Laplacians are then defined as the difference between the degree and adjacency matrices:

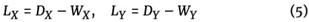

To regularize the projected representations 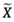 and 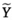,we introduce smoothness terms based on the Laplacian quadratic form. Specifically, the smoothness term for 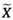 is given by:

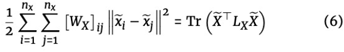

where 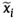 denotes the *i*-th row of 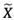. This expression encourages neighboring points in the original data space to have similar representations in the latent space, promoting smoothness in the mappings.

### GROTIA Algorithm

To integrate the datasets *X* and *Y* into a shared latent space, we propose the Graph-Regularized Optimal Transport (GROTIA) algorithm. Our objective is to find the mapping *f*_*X*_ : 𝒳 → *R*^*k*^ and *f*_*Y*_ : *y* → *R*^*k*^, that maps the data into a common *k*-dimensional space, effectively aligning their underlying manifold structures. The existence of such mapping is guaranteed by the representer theorem, which states that the optimal mappings can be expressed as finite linear combinations of the kernel functions:

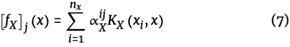

where 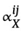 are the learned coefficients, *K*_*X*_ is the kernel function, and *x*_*i*_ are the samples from *X*. The coefficients 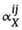 are then organized into the matrix 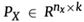, and similarity for Y we have 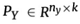. Thus, the final mapped representations are given by:

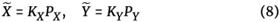

The optimization problem for the GROTIA algorithm is formulated as:

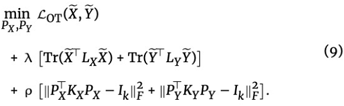

The first term 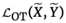 aligns the global distributions of the datasets in the latent space using sinkhorn divergence. Minimizing the Sinkhom divergence between the latent space representations ensures that the overall structures of *X* and *Y* are closely matched after projection. This captures global structural similarities and facilitates the discovery of shared manifold features between the datasets.

The second term 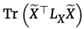 and 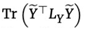 is used preserve the local geometric structures inherent in each dataset. These terms penalize the weighted differences between neighboring points in the latent space. Doing so encourages neighboring points in the ℋ_*X*,_ℋ_*X*_ to remain close in the latent space.

To prevent degenerate solutions and ensure that the mappings retain meaningful structure, we impose orthogonality constraints on the mapping matrices through the terms 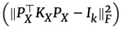 and 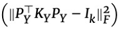.

By jointly optimizing this objective function, the GROTIA algorithm effectively balances global alignment and local structure preservation while ensuring that the projections are meaningful and well-behaved.

### Interpretable Embeddings of GROTIA

In our approach, each domain (scRNA or scATAC) is mapped into a shared, low-dimensional space using kernel-based transformations. Take the scRNA space for example, let 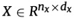 denote the data matrix for one domain, where *d*_*x*_ is the number of cells and *d*_*x*_ is the number of features. We construct a radial basis function (RBF) kernel 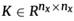, with elements 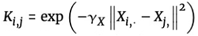 where *γ*_*X*_ is bandwidth parameter. Through our optimization procedure, we learn a coefficients matrix 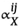 such that the *k*-dimensional embedding for thej-th cell is given by 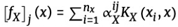. Here, if [*f*_*X*_]_*j*_ (*x*) represents the coordinate of cell *j* in the *k* dimensional learned embedding. An analogous formulation with *β* is employed for the scATAC domain (using its own kernel matrix).

To identify which original features (e.g., genes in scRNA, peaks in scATAC) have the greatest influence on each embedding dimension, we compute partialderivatives of *f*_*d*_ (*x*_*j*_) with respect to each feature. Concretely, let *x*_*j,g*_ denote the value of feature *g* in cell *j*. Then, for an RBF kernel

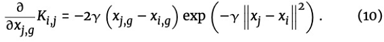

Using the chain rule, the partial derivative of the *d*-th embedding coordinate with respect to *x*_*j,g*_ becomes

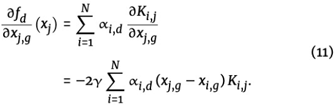

Thus, a large magnitude of 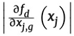 indicates that small per-turbations in feature *g* for cell *j* induce substantial shifts in the *d*-th embedding coordinate. To obtain a global feature-importance measure, we average these derivatives across all cells:

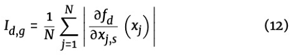

Features *g* with higher *I*_*d,g*_ are deemed more influential in shaping dimension *d*. An identical procedure is applied in the scATAC domain using the learned projection β and its kernel *K*_*Y*_.

To identify key molecular drivers in each latent dimension, we first ranked genes by their contribution scores *I*_*d,g*_. The highest-ranked genes displayed significant variation in expression closely linked to the biological structure observed within the low-dimensional embedding. Gene ontology (GO) enrichment analyses conducted on these top-ranking genes using g:Profiler [20] with default parameters revealed strong enrichment for cellular metabolism-related processes.

In parallel, we investigated regulatory elements in the chromatin accessibility (ATAC) domain. Similar to the gene-ranking procedure, open chromatin peaks were ordered by their respective contribution scores for each GROTIA-derived dimension. We then performed de novo motif discovery on the most influential peaks using GimmeMotifs [21], applying a false discovery rate (FDR) threshold of <0.001. Each discovered motif was mapped to its nearest genes within a 20 kb window around their transcription start sites (TSS), a range widely used to capture most enhancer/promoter interactions [22, 23]. This strategy identified candidate transcription factor (TF)-gene pairs whose regulatory relationships may underlie the dimension-specific chromatin states observed in the low-dimensional embedding.

### Co-Cluster Using Optimal Transport Plan

GROTIA also enables a post-integration analysis that yields data-driven clusters. Specifically, an optimal transport plan is first computed to quantify the cost or flow between two distinct sets of entities (e.g., cells in RNA and ATAC spaces). We then apply a co-clustering algorithm [24] directly to the resulting transport matrix to simultaneously group row and column entities. Treating the OT plan as a bipartite graph, the co-clustering approach identifies latent structural patterns, minimizing within-group transport costs while maximizing between-group separations.

### Evaluated metrics

Each alignment method was assessed in two distinct evaluation modes, each paired with two quantitative metrics. Unsupervised mode tuned its hyper-parameters solely by minimizing the model’s objective function, deliberately withholding any label information. Semi-supervised mode, in contrast, selected hyper-parameters that maximized downstream cell-type classification accuracy on a held-out validation set; crucially, these labels were used only during the tuning phase and were never provided to the model as inputs, preserving the semi-supervised setup.

The first metric is Fraction of Samples Closer Than the True Match (FOSCTTM). For each sample *x*_*i*_ in domain *X*, let the corresponding (true matched) sample in domain *Y* be 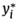. We first embed both *X* and *Y* into a common space via embedding functions *f*_1_ and *f*_2_, respectively. We then define the distance between embedded points using a distance measure *d*(·, ·). The FOSCTTM metric for each sample *x*_*i*_ measures the fraction of samples in *Y* that are closer to *x*_*i*_ (in the embedded space) than its true matchy 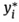. Formally,

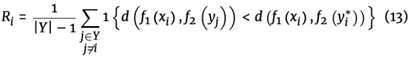

where **1**{·} is the indicator function, returning 1 if the condition is satisfied and 0 otherwise. The FOSCTTM score for the entire dataset is the average of *R*_*i*_ across all *x*_*i*_ ∈ *X* :

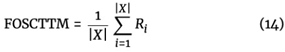

A lower FOSCTTM value indicates better alignment, as it means fewer samples in *Y* are closer to *x*_*i*_ than the true match 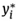.

The second metric is Label transfer accuracy (LTA) and it evaluates how well cell-type (or other categorical) labels can be transferred from domain *X* to domain *Y* in the integrated space. After embedding both datasets into a shared representation, each point *x*_*i*_ ∈ *X* has a known label *L*_*X*_ (*x*_*i*_). We define kNN (*x*_*i*_) as the set of the *k* nearest neighbors of *x*_*i*_ in the embedded representation of *Y*. Let

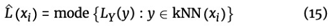

where mode (·) returns the most common label among those *k* neighbors. The LTA score is then computed as:

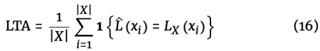

where **1**(·) is the indicator function. A higher LTA indicates that the integrated embedding preserves biological labels more accurately between the two domains.

### Training Details

We implemented GROTIA in PyTorch, using the Adam optimizer with a learning rate of 0.0001. Whenever the training loss plateaued, the learning rate was reduced by a factor of 0.5. We selected the latent dimension to be either 5 or 8 and observed that GROTIA remained robust to this choice. To preserve local structure via the graph Laplacian, we set the number of nearest neighbors to 5. Two hyperparameters appear in the loss function: *λ*, which controls distribution matching between modalities, and *ρ*, which emphasizes local geometry within each modality. We searched over a grid of *λ* ∈ { 1, 10^−1^, 10^−2^,10^−3^} and *ρ* ∈ {10^−3^, 10^−4^, 10^−5^, 10^−6^}, with the constraint *λ* > ρ to ensure that local structure is preserved and the mapping remains close to a projection.

Although GROTIA does not require identical features or genes across different modalities, it relies on the assumption of a shared underlying biology (e.g., cells of common lineages) so that datasets can be meaningfully aligned. As a standard preprocessing step to remove noisy cells and genes, we applied the following procedure to scRNA-seq data: (1) filter out cells containing fewer than 200 detected genes, (2) remove genes found in fewer than three cells, (3) log-normalize expression values by scaling each cell’s total expression to 10,000, (4) identify the 2,000 most highly variable genes, and (5) use the first 50 principal components of the processed data. For scATAC-seq data, we used a similar approach: (1) filter out cells with fewer than 200 detected genes, (2) remove genes present in fewer than three cells, (3) apply TF-IDF normalization, and (4) use the first 50 principal components.

### Baseline Settings

To benchmark our method (GROTIA), we compared it against several existing approaches, each downloaded and configured according to the authors’ guidelines. SCOT (v1.0) was obtained from https://github.com/rsinghlab/SCOT. We provided the same PCA-preprocessed input to SCOT as to GROTIA, mirroring SCOT’s original publication. We then tuned the hyperparameter based on the recommendations in the SCOT documentation.

We downloaded UnionCom (v0.4.0) and applied the same input as in GROTIA, again following the developers’ suggested preprocessing steps. All hyperparameters were tuned according to the guidance provided in the UnionCom package.

For UniPort (V1.3), which supports diagonal integration (i.e., mode=d) to align datasets without common genes, we used 2,000 highly variable genes from the scRNA – seq data and peaks exceeding a threshold ofl for the scATAC-seq data. We also employed TF-IDF normalization, replicating the tutorial steps outlined by the UniPort authors. We obtained the MMD-MA (V1.0) PyTorch implementa-tion from https://bitbucket.org/noblelab/2020_mmdma_pytorch/src. As with GROTIA and SCOT, we provided PCA-preprocessed data and tuned its hyperparameters in accordance with the authors’ guidelines.

Lastly, we downloaded scConfluence (v0.1.1). For the version without prior information, we set λ_*IOT*_ = 0, forcing a diagonal integration approach.When using scConfluence with prior information, we followed the recommended settings from the authors. We also tuned the remaining hyperparameters according to their instructions, applying identical preprocessing to ensure fair comparisons across all methods.

### Use of Large-Language Models

We used OpenAI ChatGPT only for grammar, spelling, and stylistic refinement of the manuscript. The model did not generate or alter any scientific content, data analysis, interpretations, or conclusions. All AI suggestions were manually reviewed and either accepted or rejected by the authors.

## Results

### GROTIA integrated simulated datasets in both semi and unsuperviseed setting

We evaluated the GROTIA algorithm using four simulated datasets previously discussed. Performance of the GROTIA algorithm was compared against five benchmarked methods: SCOT, Unioncom, MMD-MA, and two VAE-based approaches, Scconfluence and Uniport. Evaluations were conducted under two scenarios: semi-supervised (partial label information available) and unsupervised (no label information available).

In Figure 2a, the left column (solid color) and right column (vertical axis) display FOSCTTM and label transfer accuracy, respectively, under the semi-supervised setting. In the semi-supervised scenario, GROTIA and SCOT demonstrated consistently high performance across all datasets, with Unioncom and MMD-MA closely following. Scconfluence and Uniport showed comparatively lower performance, possibly due to their reliance on cross-modality guidance, which was not available in these simulations. In Figure 2b, each dataset from each domain is shown prior to integration, while Figure 2c shows the integrated dataset, colored by cell type on the left and by domain on the right.

**Figure 2.**
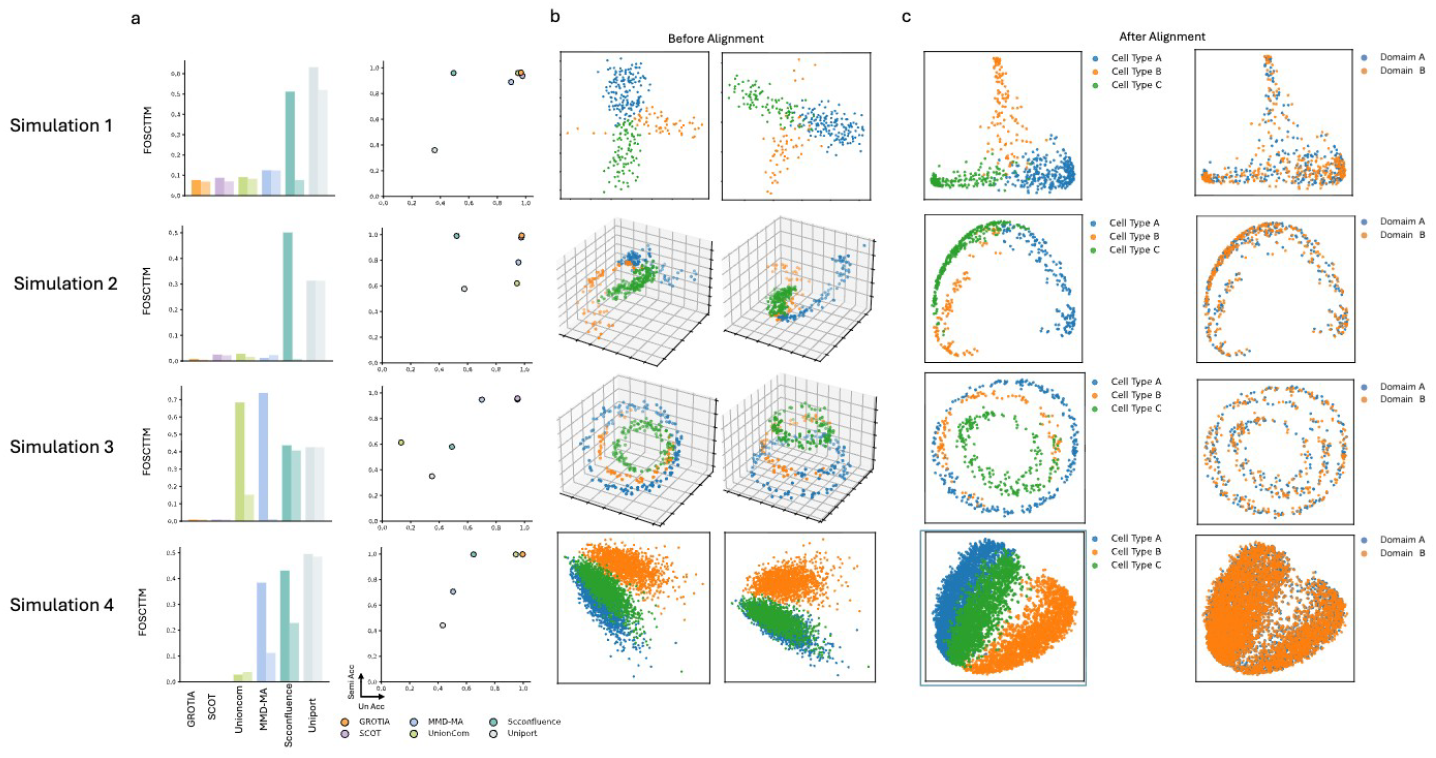
Benchmarking results on simulated datasets. a) Evaluation of Label Transfer Accuracy and Fraction of Samples Closer Than the True Match (FOSCTTM) across five benchmarked methods under two evaluation modes: semi-supervised and unsupervised. For each method, two bars are shown in the same plot, with the left bar representing the semi-supervised mode and the right bar representing the unsupervised mode for FOSCTTM. Label Transfer Accuracy is shown with semi-supervised results on the x axis and unsupervised results on the y-axis. b) Visualization of simulated datasets before integration. Simulations 1 and 4 are visualized using the first two PCA components, whereas Simulations 2 and 3 are visualized using the first three axes from multidimensional scaling (MDS). c) Visualization after integration. The first columndisplays data colored by cell type, while the second column shows data colored by domain. All four datasets are visualized using the first two PCA components.

In Figure 2a, the left column (solid color) and right column (vertical axis) display FOSCTTM and label transfer accuracy, respectively, under the semi-supervised setting. In the unsupervised scenario, where alignment depends solely on intrinsic structural information, GROTIA and SCOT maintained relatively stable performance. Other methods showed varying degrees of accuracy reduction, notably Unioncom on dataset 3, MMD-MA on datasets 3 and 4, Scconfluence on datasets 1, 2, and 4, and Uniport on dataset 1. These variations highlight the challenges inherent in unsupervised alignment without label guidance.

The stable performance of GROTIA can be attributed to their effective use ofWasserstein-based (WD) losses, capturing intrinsic geometric structures. Additionally, GROTIA employs orthogonality constraints within the Reproducing Kernel Hilbert Space (RKHS), enhancing embedding interpretability and stability. Although VAE-based methods also utilize WD losses, their embeddings can be susceptible to rotational variations, affecting alignment stability in the absence of label information. Detailed numerical results corresponding to the metric plots are presented in Supplementary Tables A2-A5.

### GROTIA integrated real word datasets in both semi and unsupervised setting

We then evaluated the GROTIA algorithm on four real-world datasets.We chose these paired multi-omics datasets specifically to enable diagonal integration with known ground-truth cell correspondences. Importantly, during benchmarking, all methods are provided with unpaired data, and the known cell-pairing information was used only for evaluating alignment accuracy.

The benchmarking process used here follows the same procedure described in the simulation study, with one modification: for the PBMC-1 and PBMC-2 datasets, we additionally include scConfluence with prior information. We do not include the prior version of scConfluence for the scGEM and SNARE datasets because the original authors performed dimensionality reduction to process these data, meaning commonly used gene selection methods are not available and the resulting feature space is small.

In Figure 3a, the left column (solid color) and right column (vertical axis) display FOSCTTM and label transfer accuracy, respectively, under the semi-supervised setting. Across the scGEM and SNARE-seq datasets, GROTIA achieves the second-best FOSCTTM in scGEM and the best in SNARE-seq, while attaining the highest label transfer accuracy in both. SCOT obtains the top FOSCTTM in scGEM but ranks second in the remaining evaluations. scConfluence (default), MMD-MA, UnionCom, and UniPort follow in overall performance. For the two PBMC datasets, GROTIA places second in FOSCTTM for both PBMC-1 and PBMC-2; in PBMC-2, scConfluence with prior information attains the best FOSCTTM. Regarding label transfer accuracy, GROTIA ranks second in PBMC-1 and first in PBMC-2. In Figure 3b, each dataset from each domain is shown prior to integration, while Figure 3c shows the integrated dataset, colored by cell type on the left and by domain on the right.

**Figure 3.**
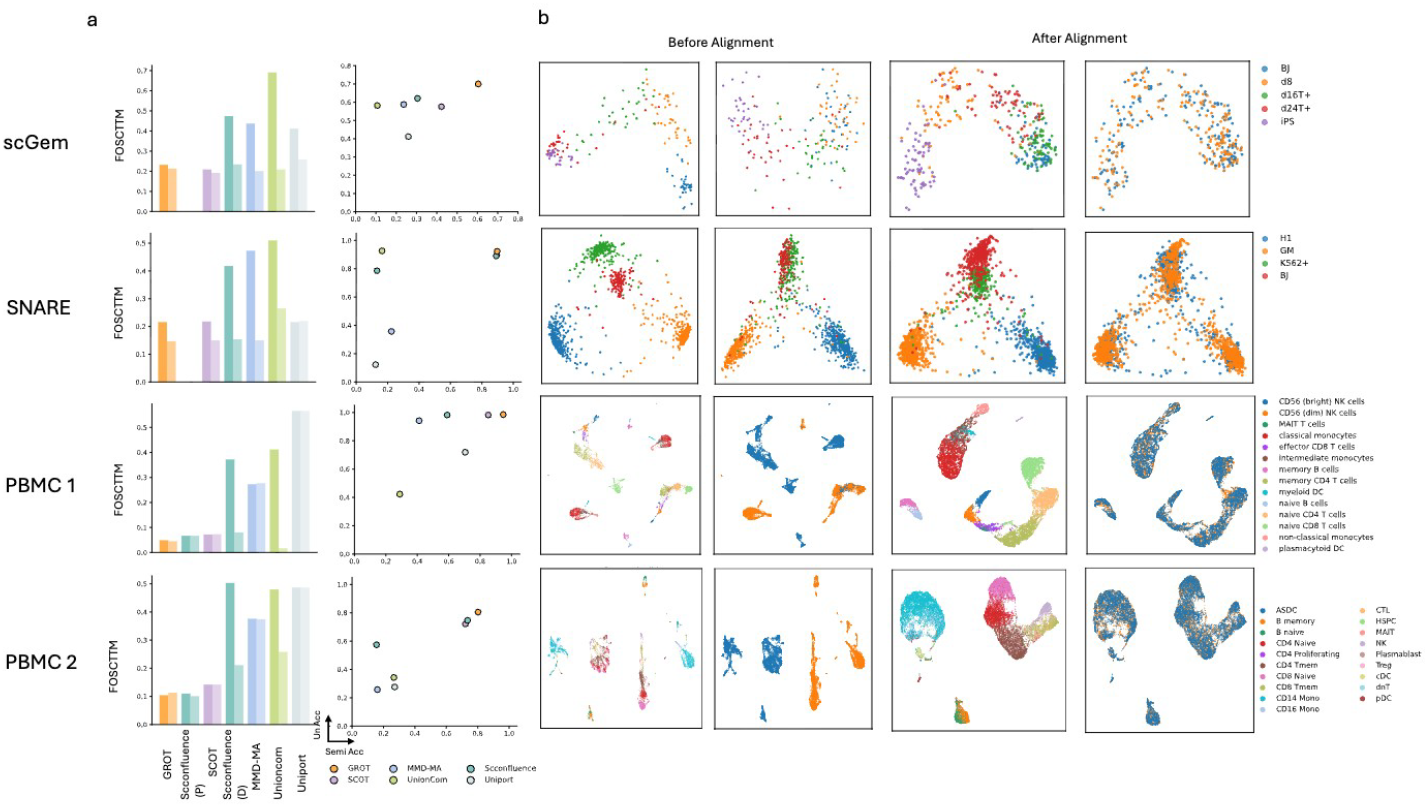
Benchmarking results on real word datasets. a) Evaluation of Label Transfer Accuracy and Fraction of Samples Closer Than the True Match (FOSCTTM) across five benchmarked methods under two evaluation modes: semi-supervised and unsupervised. For each method, two bars are shown in the same plot, with the left bar representing the semi-supervised mode and the right bar representing the unsupervised mode for FOSCTTM. Label Transfer Accuracy is shown with semi-supervised results on the x-axis and unsupervised results on the y-axis. b) Visualization of real word datasets before integration. ScGem and SNARE are visualized using the first two PCA components, whereas PBMC 1 and PBMC 2 are visualized using the first two UMAP Components. c) Visualization alter integration. The first column displays data colored by cell type, while the second column shows data colored by domain. ScGem and SNARE are visualized using the first two PCA components, whereas PBMC 1 and PBMC 2 are visualized using the first two UMAP Components.

Under the fully unsupervised setting, shown by lighter colors in the left column of Figure 3a (and the right column’s horizontal axis for label transfer accuracy), GROTIA demonstrates the strongest overall performance in both FOSCTTM and label transfer accuracy across all four real-world datasets, except for ranking second in FOSCTTM on scGEM. SCOT and scConfluence exhibit comparable results, followed by UnionCom, MMD-MA, and UniPort. Detailed numerical results corresponding to the metric plots are presented in Supplementary Tables A6-A9.

### GROTIA Reveals Gene-Specific Contributions and Key Biological Processes in the RNA Embedding

GROTIA provides an in-model measure of gene importance, pin-pointing which genes drive variation along each latent dimension. Specifically, we compute partial derivatives of the RBF kernel embeddings with respect to each gene’s expression and then weight these by the projection matrix to obtain a contribution score for every gene–dimension pair. Ranking these scores identifies dimension-specific “signature genes”–those whose expression changes most strongly reposition cells in the low-dimensional space. For additional details, see Section.

Figure 4a displays the overall gene importance scores across all eight GROTIA-derived dimensions. Figure 4b presents UMAP visualizations of the top gene expression patterns for Dimensions 1 and 3, illustrating how high-impact genes are distributed across the dataset. For Dimension 1, the top three contributing genes (LYZ, ZEB2, PLXDC2) are highly expressed in monocytes and myeloid cell populations, consistent with further differentiation within the monocyte lineage. Moreover, Dimension 3 emphasizes GNLY, CCL5, and LEFl – genes enriched in NK cells and T cells, indicating a T cell–specific transcriptional program. Notably, GROTIA requires no a priori matching of features across modalities, so these dimension-specific drivers offer an unbiased method to uncover potential marker genes.

**Figure 4.**
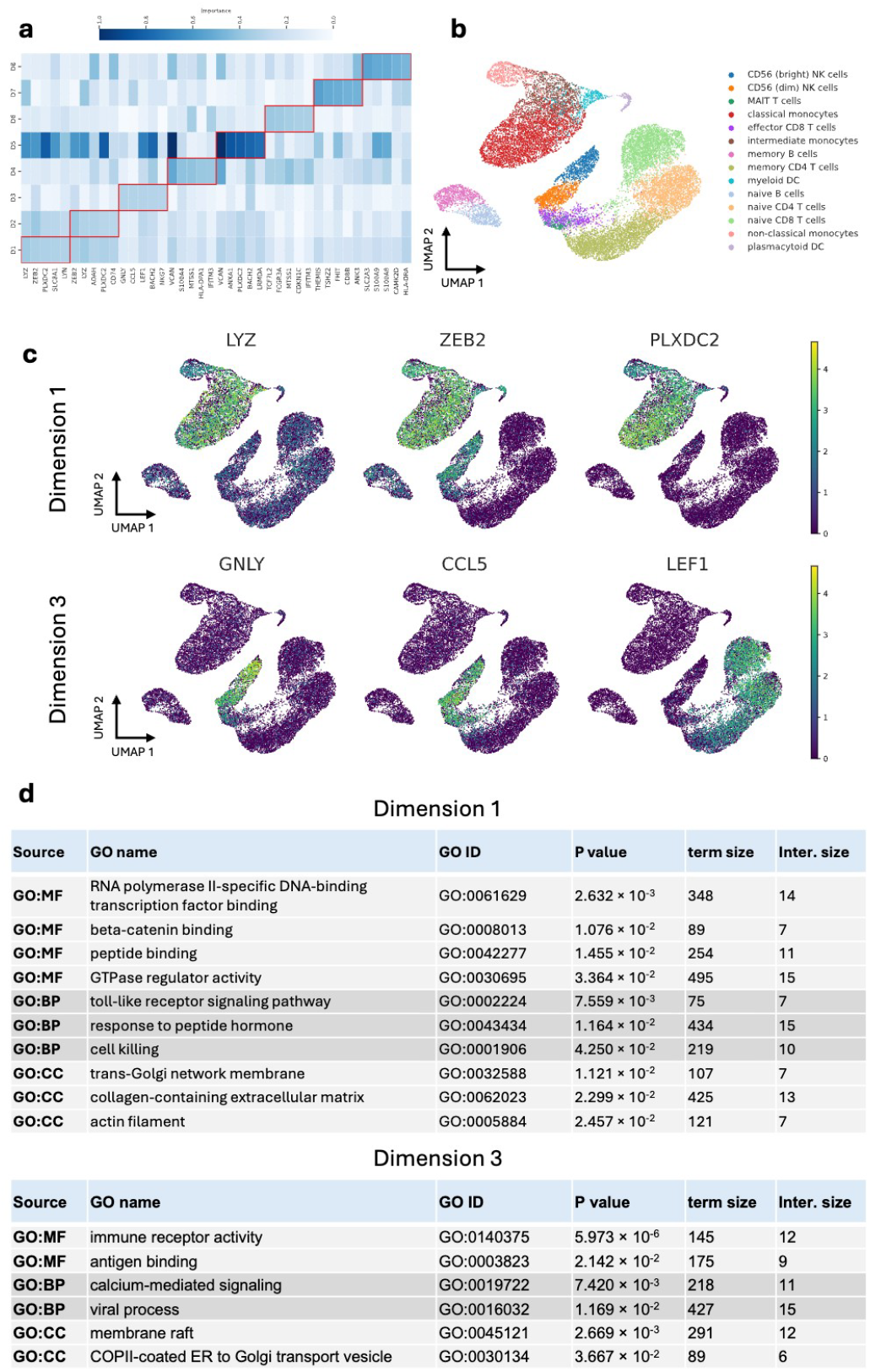
(a) Heatmap of partial derivative based importance scores for the top five genes in each of the eight GROTIA derived dimensions (D1–D8). Darker blue indicates higher importance. The top five genes per dimension are outlined in the red box. (b) UMAP projections of Dimension 1 and Dimension 3, highlighting cell-type annotations (top) and the expression distributions of top-ranked genes (middle and bottom). Warmer hues denote higher expression, revealing distinct cellular subsets for Dimension 1 and Dimension 3. (c) Summaries from the Gene Ontology experiment, showing significant enrichment grouped by Molecular Function (MF), Cellular Component (CC), and Biological Process (BP). Collectively, these panels show how GROTIA’s dimension-wise interpretability links high-impact genes to key biological functions.

To link the top contributors in Dimension 1 to biological processes, we performed Gene Ontology (GO) enrichment analysis, as shown in figure Fig. 4c. These genes were enriched for the terms RNA polymerase II-specific DNA-binding transcription factor binding, β-catenin binding, and peptide binding. The first term suggests that Dimension 1 captures a transcriptional regulatory program active during monocyte and dendritic cell differentiation, involving lineage-defining regulators such as ZEB2, which is essential for monocyte and plasmacytoid DC development [25]. The second highest-ranked term, β-cateninbinding, indicates involvement of Wnt/ β-catenin signaling in monocyte and DC biology; for instance, β-catenin activation fosters a tolerogenic phenotype in DCs [26], while its aberrant stabilization can obstruct normal monocyte-macrophage differentiation [27]. Finally, enrichment for peptide binding aligns with the antigen processing and presentation roles of monocytes and DCs, consistent with elevated HLA-DR expression in intermediate monocytes [28]. For additional UMAP distributional plots of top genes, see Supplementary Figures A.1 and A.2.

In Dimension 3 of the PBMC transcriptional analysis, we observed GO term enrichment related to immune receptor activity, antigen binding, and calcium-mediated signaling. This suggests that Dimension 3 captures variation in lymphocyte receptor expression and signaling. The GO category immune receptor activity is associated primarily with T lymphocytes, which uniquely express the T-cell receptor complex (for example, CD3 subunits and TCR *α/β* chains) mediating antigen-specific recognition [29]. Moreover, antigen binding reflects the high expression of immunoglobulin genes by B cells, consistent with their exclusive role in producing antigen-specific antibodies [30]. Finally, calcium-mediated signaling highlights a key activation pathway in T cells, where antigen-receptor engagement triggers *Ca*^2+^ influx through store-operated *Ca*^2+^ channels to activate downstream effectors such as the calcineurin-NFAT pathway, a process essential for lymphocyte activation [31]. For additional Gene Ontology enrichment results corresponding to the remaining dimensions, see Supplementary Tables A10–A15.

### GROTIA Identifies High-Impact Peaks and Regulatory Mechanisms in the ATAC Embedding

Similarly, in the ATAC domain, we ranked open chromatin peaks by their gradient-based contribution scores and performed motif discovery on the highest-impact peaks (see Methods). Mapping each motif to its nearest gene within a 20 kb window revealed putative regulatory relationships linking epigenetic accessibility to transcriptional output. Figure 5a illustrates the procedure for identifying transcription factor–gene pairs from the top-ranked peaks. For further details, please refer to Section.

**Figure 5.**
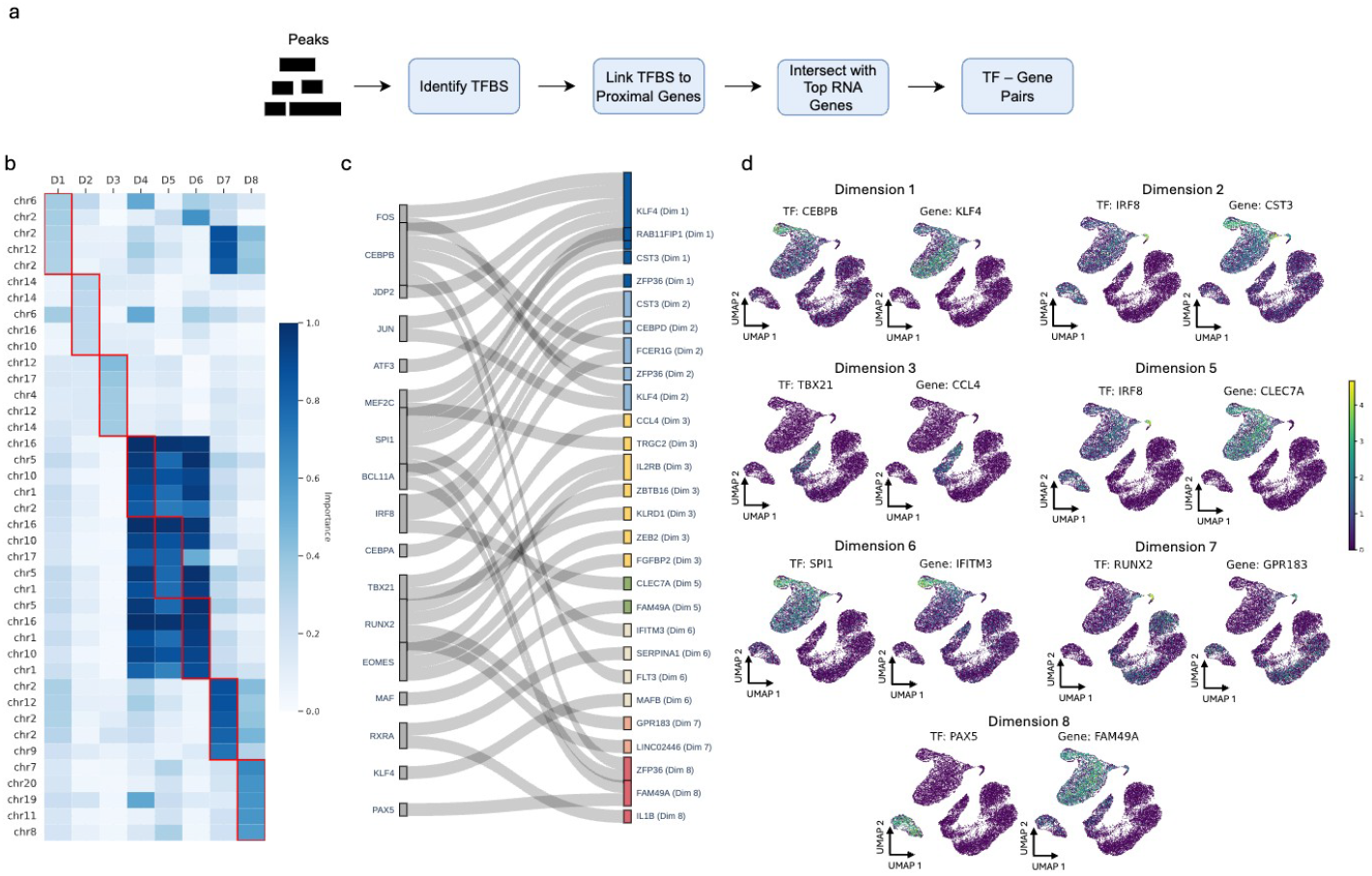
(a) Schematic of the workflow for identifying significant motifs in open chromatin peaks, mapping these motifs to nearby genes, and intersecting them with top RNA space genes to form motif–gene pairs. (b) Heatmap of gradient-based importance scores for the five most influential ATAC peaks in each GROTIA-derived dimension (D1–D8), where darker blue indicates greater importance. (c) Sankey diagram illustrating dimension-specific gene–factor pairs. Genes (left) connect to their putative transcription factors (right), validated through literature. For instance, FOS are implicated as potential regulators of Gene KFL4 in Dimensions 1and CEBPB as potential regulators of Gene FCR1G in dimension 2. (d) UMAP embeddings colored by accessibility values of selected gene–factor pairsfrom Dimensions 1, 3, 5, 6, 7, and 8. Warmer hues denote higher accessibility, highlighting dimension-specific regulatory landscapes. Co-expression patterns further support these putative regulatory relationships.

Figure 5b highlights the top-ranked peaks (based on partial derivative-based importance) across the eight GROTIA-derived dimensions (D1–D8), with the most influential peaks in each dimension outlined in red boxes. Figure 5c shows the inferred transcription factor – gene pairs associated with each dimension. Finally, Figure 5d offers a UMAP projection of these pairs, visually illustrating potential co-expression or repression relationships. In dimension 1, we identified a CEBPB– KLF4 pair. CEBPB is essential for proper monocyte development, including the survival of certain subsets, and likely induces KLF4 as part of the monocyte differentiation network. Indeed, PU.1 (encoded by SPI1) directly upregulates KLF4, and CEBPB cooperates with PU.1 to drive monopoiesis [32]. In dimension 2, the IRF8 –CST3 pair emerged. IRF8 directly activates CST3 (cystatin C) during macrophage differentiation, mediated by a unique promoter element that overlaps IRF and ETS sites and requires both IRF8 and an ETS partner (e.g., PU.1) [33]. Dimension 3 highlighted TBX21 (T-bet) –CCL4, wherein T-bet binds to and positively regulates CCL4 in Th1 cells. Genome-wide ChIP-chip experiments in human T cells confirmed CCL4 as a direct T-bet target, revealing T-bet binding in regulatory regions that activate CCL4 transcription [34]. In dimension 5, we found IRF8–CLEC7A. Genome-wide binding studies have identified CLEC7A as an IRF8 target in human myeloid cells [35], and co-expression networks further link CLEC7A with an IRF8-centered module. Consistently, aging human microglia upregulate CLEC7A alongside other “activated” microglial genes under the control of an IRF8/SPI1/RUNX1/TAL1 network [36]. In dimension 6, the SPI1–IFITM3 pair indicates that PU.1 controls a broad antiviral gene program in macrophages, with IFITM3 explicitly citetd as a PU.1-regulated antiviral factor [37]. Dimension 7 highlighted RUNX2–GPR183-Finally, in dimension 8, the PAX5–FAM49A pair was supported by evidence of PAX5 ChIP-seq peaks near FAM49A [38]. Notably, FAM49Ais expressed at lower levels in PAX5-positive pro–B cells and is de-repressed in PAX5-deficient cells, indicating that PAX5 normally suppresses Fam49a expression during early B-cell development [39].

### GROTIA enables identification of cellular subpopulation on integrated space

Beyond simply projecting scRNA and scATAC profiles into a shared space, GROTIA provides a natural framework for discovering sub-populations in an unsupervised manner. While many datasets come with predefined labels (e.g., annotated cell types), these annotations may be incomplete or coarse, especially when new subtleties or states exist that were not recognized during initial labeling. By clustering cells within GROTIA’s integrated representation, we can uncover finer structures and novel cellular states that may otherwise remain masked by legacy annotations.

To determine the optimal number of clusters *k*, we plot the reconstruction error for various *k*-values and look for a distinct elbow (Fig. 6a). Beyond this point, further increasing *k* offers minimal improvement in accuracy while potentially fragmenting biologically coherent groups. We therefore select the *k* at the bump, yielding a robust trade-off between clustering granularity and data fidelity. We then visualize the resulting assignments alongside the original cell-type labels on a UMAP projection (Fig. 6c). Notably, GROTIA identifies distinct subpopulations that align well with known major cell types, yet can also isolate refined subclusters reflecting subtle transcriptional and epigenetic differences.

**Figure 6.**
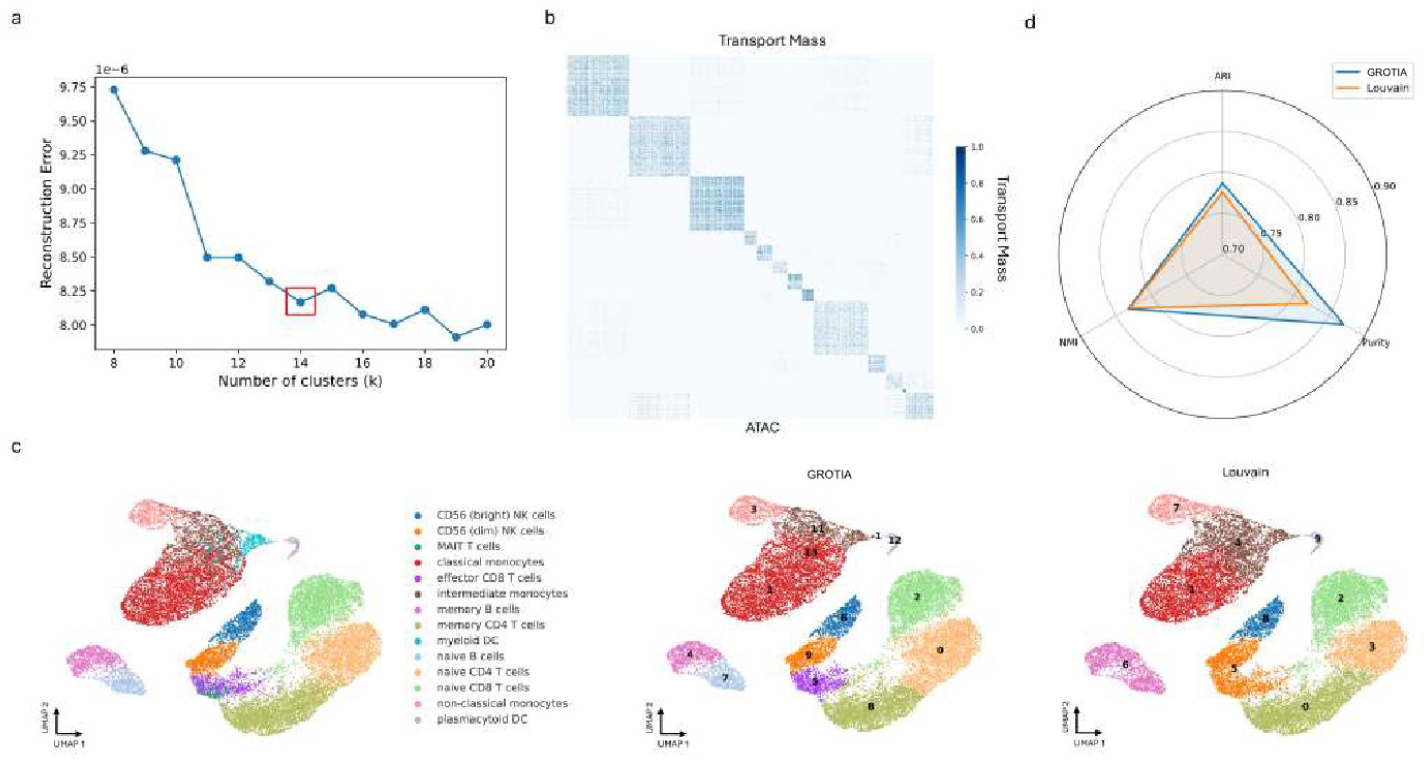
(a) Reconstruction error plotted against the number of clusters k, with the chosen k marked by a red box. This optimal k balances clustering granularity and data fidelity. (b) Heatmap of the transport mass after co-clustering, illustrating how GROTIA aligns cells from scRNA and scATAC. The block-diagonal pattern indicates coherent groupings across both modalities. (c) UMAP projections of the integrated dataset, colored by true cell type annotations (left), and clustered with GROTIA (center) or Louvain (right). For each method, predictedclusters are labeled by the ground-truth cluster with which they most overlap. GROTIAproduces distinct subpopulations consistent with known cell type boundaries. (d) Radar plot comparing GROTIA (orange) and Louvain (blue) on three clustering metrics (ARI, NM!, and Purity). GROTIA demonstrates higher or comparable performance, indicating its ability to robustly identify meaningful subpopulations.

Finally, to benchmark the quality of GROTIA’s partitioning, we compare against a conventional community detection algorithm (Louvain) using three standard clustering metrics: Adjusted Rand Index (ARI), Normalized Mutual Information (NMI), and Purity (Fig. 6d). Our method achieves comparable or better performance, demonstrating that GROTIA’s alignment-based approach not only reconciles multi-omic data but also preserves biologically meaningful structures when clustering.

## Discussion

We present GROTIA, a novel algorithm for unsupervised single-cell multi-omics integration that combines optimal transport for global alignment with graph regularization to preserve local structure. Benchmarking against state-of-the-art methods in both unsupervised and semi-supervised settings, GROTIA delivers comparable or superior performance while offering a computationally efficient solution. Critically, our framework also includes an in-model interpretability mechanism, allowing users to identify which genes or peaks drive each dimension of the integrated embedding. This enables targeted downstream analyses–such as Gene Ontology enrichment or motif discovery–to reveal meaningful biological processes in the RNA and ATAC spaces.

Beyond aligning multi-omics data, GROTIA leverages its transport plan for post-integration clustering, offering a data-driven approach to refine or correct misannotations in existing labels. Notably, unlike methods that require shared features across modalities, GROTIA only assumes that cells (rather than individual genes or peaks) follow a similar distribution if they belong to the same type or lineage—thus broadening its applicability to complex datasets.

Looking ahead, we plan to extend GROTIA to time-series multi-omics data, where paired measurements across multiple time points are becoming increasingly common. Furthermore, we will deepen our driver-gene analyses to decode how specific features shape the integrated embedding, with the goal of uncovering more nuanced regulatory processes across different cell states.

## Availability of source code and requirements (optional, if code is present)

Lists the following:

- Project name: GROTIA
- Project home page: https://github.com/PennShenLab/GROTIA
- Operating system(s): Platform independent Programming language: Python 3.8 or higher
- License: License: MIT License

This needs to be under an Open Source Initiative approved license where practicable compiled running software is made available. If the code is not hosted in a repository the *GigaScience* GitHub repository is also available for this purpose.

## Data Availability

The GROTIA algorithm is freely available at https://github.com/PennShenLab/GROTIA. All data used in this manuscript is publicly available and can be found at Liu et al. [11], Cheow et al. [16], Demetci et al. [12], Chen et al. [17], Cao et al. [14], and Samaran et al. [13].

## List of abbreviations

FOSCTTM: Fraction of Samples Closer Than the True Match
GO: Gene Ontology
GUMA: Generalized Unsupervised Manifold Alignment
GROTIA: (Graph-Regularized Optimal Transport Framework for Diagonal Single-Cell Integrative Analysis)
LTA: Label Transfer Accuracy
MMD: Maximum Mean Discrepancy
PBMC: Peripheral Blood Mononuclear Cell
SCOT: Single-Cell Multi-Omics Alignment with Optimal Transport
TF: Transcription Factor
TSS: Transcription Start Sites
UnionCom: Unsupervised Topological Alignment for Single-Cell Multi-Omics Integration
WD: Wasserstein-based

## Author contributions statement

Conceptualization, Z.W., Q.Z., M.K., and L.S.; Methodology, Z.W., Q.Z., and L.S.; Resources, L.S.; Formal analysis, Z.W., Q.Z., S.Y., Z.Z., M.K., T.Z., and L.S.; Writing-Original Draft, Z.W., Q.Z., Z.Z., and L.S.; Funding acquisition, L.S.; Writing-Review and Editing, Z.W., Q.Z., S.Y., Z.Z., M.K., T.Z., and L.S.

## Appendix

**Table A1.** Comparison of runtime performance (minutes) for benchmarked methods using the PBMC dataset (9,378 cells).

| Method | GROTIA | UniPort | scConfluence | Unioncom | SCOT | MMD-MA |
| --- | --- | --- | --- | --- | --- | --- |
| Runtime (min) | 15 | 17 | 20.44 | 65 | 185 | 470 |

**Table A2.** Alignment performance by FOSCTTM under unsupervised setting (First 4 columns: Simulation 1, Simulation 2, Simulation 3, Synthetic RNA-seq).

|  | Simulation 1 | Simulation 2 | Simulation 3 | Synthetic RNA-seq |
| --- | --- | --- | --- | --- |
| SCOT | 0.088 | 0.025 | **0.009** | 0.001 |
| MMD-MA | 0.125 | 0.012 | 0.739 | 0.384 |
| UnionCom | 0.091 | 0.028 | 0.684 | 0.028 |
| Uniport | 0.632 | 0.313 | 0.426 | 0.495 |
| Scconfluence | 0.512 | 0.501 | 0.437 | 0.431 |
| GROTIA(Ours) | **0.077** | **0.008** | **0.009** | **5e-5** |

**Table A3.** Alignment performance by FOSCTTM under unsupervised setting (Last 2 columns: scGEM and SNAREseq).

|  | scGEM | SNAREseq | pbmc10X | pmbc |
| --- | --- | --- | --- | --- |
| SCOT | **0.209** | 0.218 | 0.072 | 0.142 |
| MMD-MA | 0.437 | 0.473 | 0.273 | 0.376 |
| UnionCom | 0.691 | 0.510 | 0.412 | 0.480 |
| Uniport | 0.412 | 0.216 | 0.566 | 0.487 |
| Scconfluence(Diagonal) | 0.474 | 0.418 | 0.372 | 0.503 |
| Scconfluence(Prior) | - | - | 0.067 | 0.110 |
| GROTIA(Ours) | 0.215 | **0.216** | **0.049** | **0.104** |

**Figure A.1.**
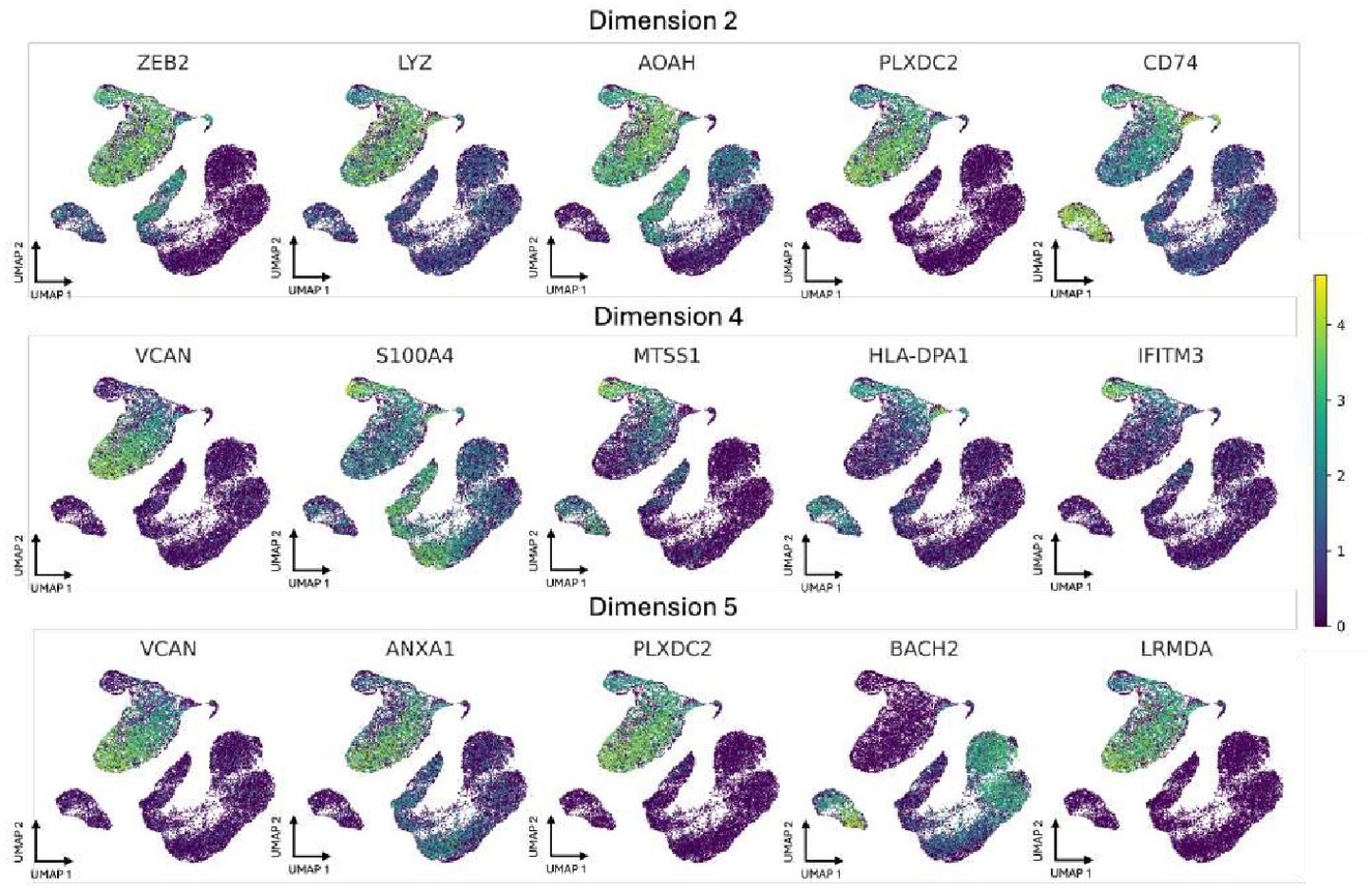
UMAP Plot of Top Genes for Dimensions 2, 4, and 5.

**Table A4.**
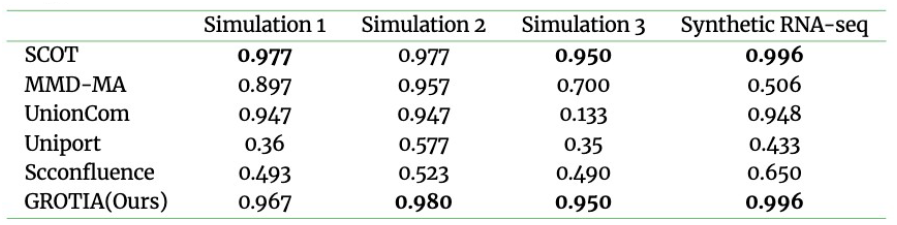
Alignment performance by label transfer accuracy (*k* = 5) under unsupervised setting (First 4 columns: Simulation 1, Simulation 2, Simulation 3, Synthetic RNA-seq).

**Table A5.** Alignment performance by label transfer accuracy (*k* = 5) under unsupervised setting (Last 2 columns: scGEM and SNAREseq).

|  | scGEM | SNAREseq | pbm10X | pmmc |
| --- | --- | --- | --- | --- |
| SCOT | 0.423 | 0.852 | 0.894 | 0.722 |
| MMD-MA | 0.237 | 0.412 | 0.224 | 0.161 |
| UnionCom | 0.107 | 0.288 | 0.164 | 0.266 |
| Uniport | 0.260 | 0.705 | 0.123 | 0.271 |
| Scconfluence(Diagonal) | 0.305 | 0.590 | 0.132 | 0.155 |
| Scconfluence(Prior) | – | – | 0.891 | 0.735 |
| GROTIA(Ours) | **0.588** | **0.947** | **0.897** | **0.802** |

**Table A6.** Alignment performance by FOSCTTM (The lower the better) under semi-supervised setting for the first four datasets.

|  | Simulation 1 | Simulation 2 | Simulation 3 | Synthetic RNA-seq |
| --- | --- | --- | --- | --- |
| SCOT | 0.070 | 0.022 | **0.009** | 0.001 |
| MMD-MA | 0.124 | 0.023 | 0.012 | 0.112 |
| UnionCom | 0.083 | 0.016 | 0.152 | 0.038 |
| Uniport | 0.520 | 0.313 | 0.426 | 0.485 |
| Scconfluence | 0.077 | 0.007 | 0.407 | 0.228 |
| GROTIA(Ours) | **0.069** | **0.005** | **0.009** | **1e-6** |

**Table A7.** Alignment performance by FOSCTTM (The lower the better) under semi-supervised setting for scGEM, SNAREseq, pbmc10X, and pbmc.

|  | scGEM | SNAREseq | pbm10X | pmmc |
| --- | --- | --- | --- | --- |
| SCOT | **0.192** | 0.150 | 0.073 | 0.142 |
| MMD-MA | 0.201 | 0.150 | 0.277 | 0.374 |
| UnionCom | 0.209 | 0.265 | **0.017** | 0.258 |
| Uniport | 0.259 | 0.220 | 0.566 | 0.487 |
| Scconfluence (Diagnola) | 0.234 | 0.154 | 0.080 | 0.211 |
| Scconfluence(Prior) | – | – | 0.067 | **0.101** |
| GROTIA(Ours) | 0.213 | **0.148** | 0.045 | 0.113 |

**Table A8.** Alignment performance by label transfer accuracy (*k* = 5) (The higher the better) under semi-supervised setting for the first four datasets.

|  | Simulation 1 | Simulation 2 | Simulation 3 | Synthetic RNA-seq |
| --- | --- | --- | --- | --- |
| SCOT | 0.937 | 0.977 | **0.957** | **0.998** |
| MMD-MA | 0.890 | 0.783 | 0.947 | 0.706 |
| UnionCom | 0.960 | 0.620 | 0.613 | 0.997 |
| Uniport | 0.360 | 0.577 | 0.350 | 0.442 |
| Scconfluence | 0.960 | 0.990 | 0.580 | 0.997 |
| GROTIA(Ours) | **0.963** | **0.993** | 0.950 | **0.998** |

**Table A9.** Alignment performance by label transfer accuracy (*k* = 5) (The higher the better) under semi-supervised setting for scGEM, SNAREseq, pbmc10X, and pbmc.

|  | scGEM | SNAREseq | pbm10X | pmmc |
| --- | --- | --- | --- | --- |
| SCOT | 0.576 | 0.982 | 0.894 | 0.722 |
| MMD-MA | 0.588 | 0.942 | 0.357 | 0.258 |
| UnionCom | 0.582 | 0.423 | **0.926** | 0.343 |
| Uniport | 0.412 | 0.719 | 0.123 | 0.276 |
| Scconfluence(Diagonal) | 0.621 | 0.982 | 0.786 | 0.574 |
| Scconfluence(Prior) | – | – | 0.891 | 0.747 |
| GROTIA(Ours) | **0.700** | **0.986** | 0.922 | **0.805** |

**Table A10.** Gene Ontology enrichment analysis for Dimension 2.

| source | GO name | GO ID | p_value | term size | inter. size |
| --- | --- | --- | --- | --- | --- |
| GO:MF | calcium-dependent protein binding | GO:0048306 | $4.566e - 4$ | 80 | 8 |
| GO:MF | GTPase regulator activity | GO:0030695 | $2.350e - 3$ | 495 | 17 |
| GO:BP | toll-like receptor signaling pathway | GO:0002224 | $8.450e - 3$ | 75 | 7 |
| GO:BP | [l]positive regulation of NF-kappaB | GO:0051092 | $2.748e - 2$ | 124 | 8 |
| transcription factor activity |  |  |  |  |  |
| GO:BP | receptor internalization | GO:0031623 | $3.464e - 2$ | 128 | 8 |
| GO:CC | membrane raft | GO:0045121 | $4.975e - 4$ | 291 | 13 |
| GO:CC | trans-Golgi network membrane | GO:0032588 | $1.304e - 3$ | 107 | 8 |
| GO:CC | collagen-containing extracellular matrix | GO:0062023 | $2.733e - 2$ | 425 | 13 |
| GO:CC | actin filament | GO:0005884 | $2.747e - 2$ | 121 | 7 |
| GO:CC | cell leading edge | GO:0031252 | $3.002e - 2$ | 429 | 13 |

**Table A11.**
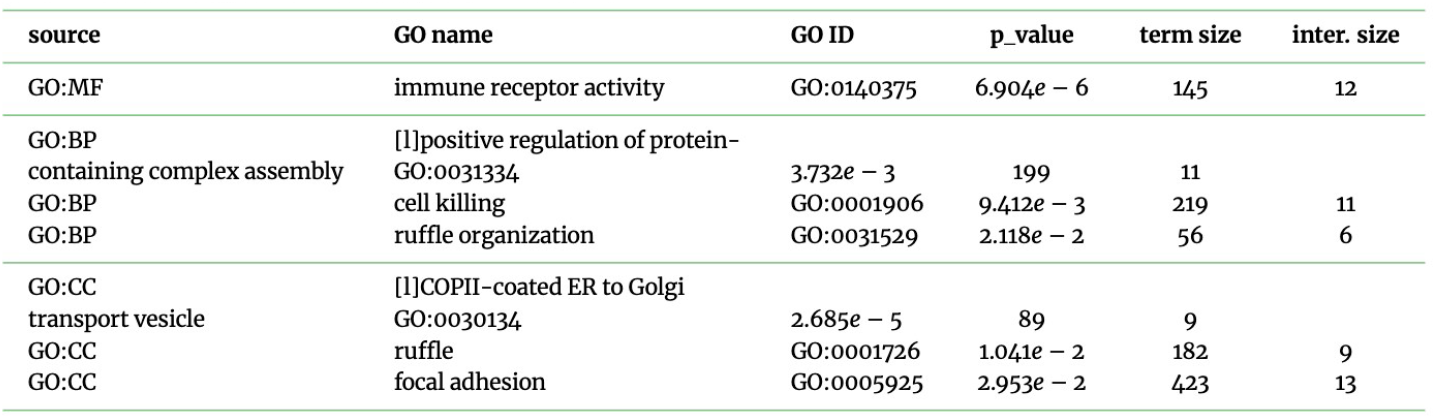
Gene Ontology enrichment analysis for Dimension 4.

**Table A12.** Gene Ontology enrichment analysis for Dimension 5.

| source | GO name | GO ID | p_value | term size | inter. size |
| --- | --- | --- | --- | --- | --- |
| GO:MF | GTPase regulator activity | GO:0030695 | $2.006e - 5$ | 495 | 20 |
| GO:MF | phospholipid binding | GO:0005543 | $1.540e - 3$ | 484 | 17 |
| GO:BP | [l]negative regulation of |  |  |  |  |
| protein phosphorylation | GO:0001933 | $1.810e - 4$ | 270 | 14 | |
| GO:BP | [l]positive regulation of NF-kappaB |  |  |  |  |
| transcription factor activity | GO:0051092 | $2.258e - 2$ | 124 | 8 | |
| GO:BP | regulation of GTPase activity | GO:0043087 | $2.717e - 2$ | 210 | 10 |

**Table A13.** Gene Ontology enrichment analysis for Dimension 6.

| source | GO name | GO ID | p_value | term size | inter. size |
| --- | --- | --- | --- | --- | --- |
| GO:MF | [l]DNA-binding transcription |  |  |  |  |
| factor binding | GO:0140297 | $3.481e - 2$ | 490 | 15 | |
| GO:MF | immune receptor activity | GO:0140375 | $3.850e - 2$ | 145 | 8 |
| GO:BP | phagocytosis | GO:0006909 | $6.492e - 6$ | 234 | 15 |
| GO:BP | cellular response to metal ion | GO:0071248 | $5.224e - 4$ | 200 | 12 |
| GO:BP | icosanoid biosynthetic process | GO:0046456 | $1.485e - 3$ | 57 | 7 |
| GO:BP | cell killing | GO:0001906 | $9.412e - 3$ | 219 | 11 |
| GO:BP | [l]positive regulation of |  |  |  |  |
| interleukin-1 beta production | GO:0032731 | $3.171e - 2$ | 60 | 6 | |

**Table A14.** Gene Ontology enrichment analysis for Dimension 7.

| source | GO name | GO ID | p_value | term size | inter. size |
| --- | --- | --- | --- | --- | --- |
| GO:MF | immune receptor activity | GO:0140375 | $3.729e - 5$ | 145 | 11 |
| GO:BP | cell-matrix adhesion | GO:0007160 | $4.671e - 2$ | 236 | 10 |
| GO:CC | T cell receptor complex | GO:0042101 | $4.214e - 3$ | 136 | 8 |

**Table A15.**
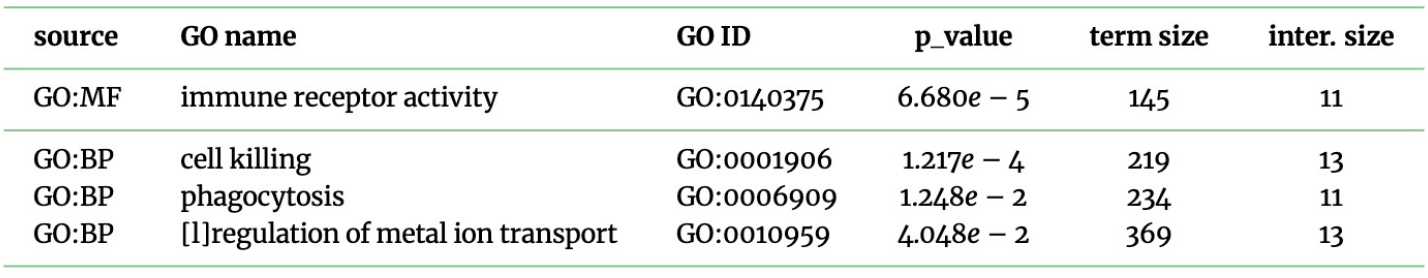
Gene Ontology enrichment analysis for Dimension 8.

**Figure A.2.**
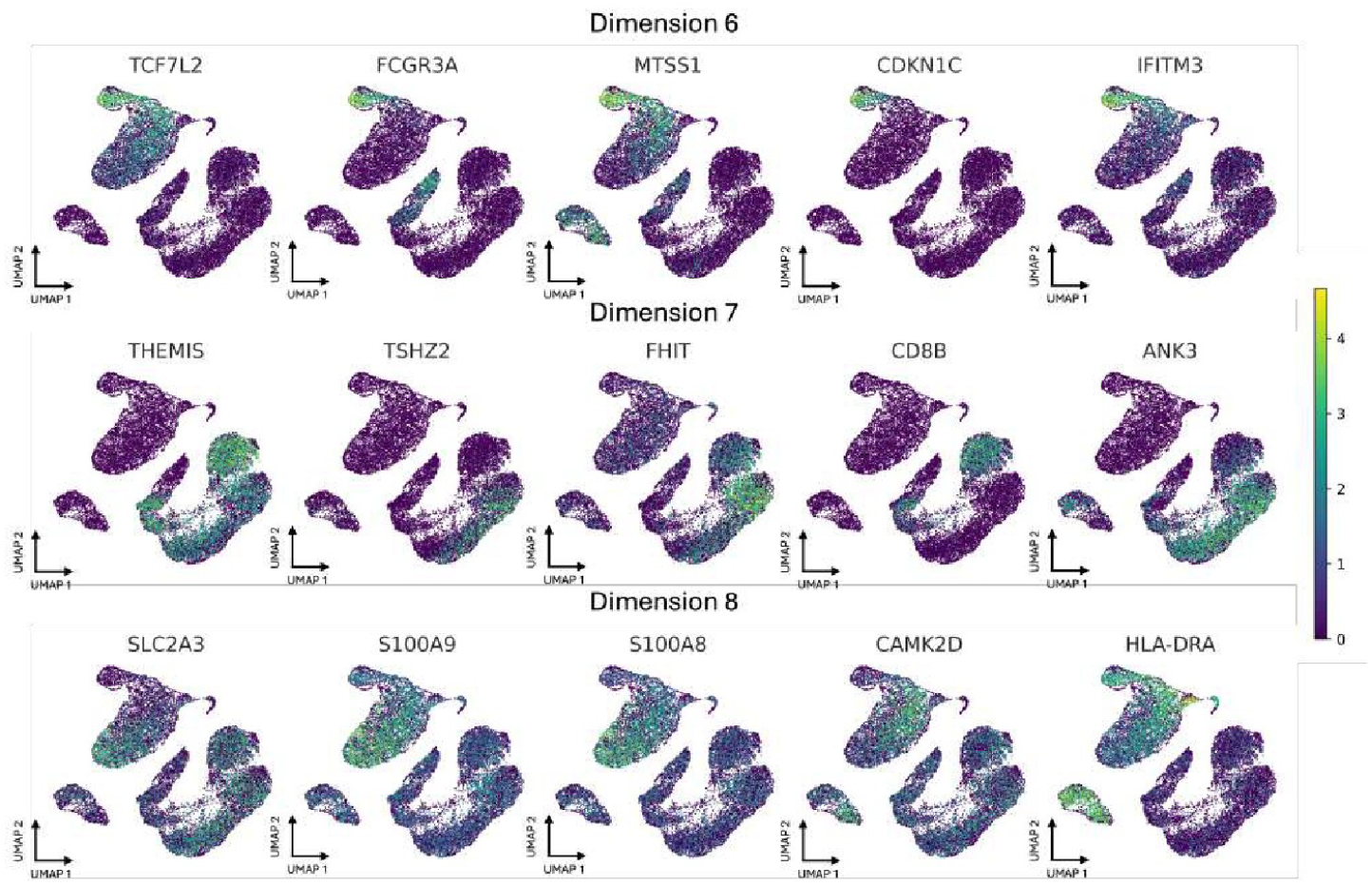
UMAP Plot of Top Genes for Dimensions 6, 7, and 8.

